# RNA-seq analyses of molecular abundance (RoMA) for detecting differential gene expression

**DOI:** 10.1101/410985

**Authors:** Guoshuai Cai, Jennifer M. Franks, Michael L. Whitfield

## Abstract

**Motivation:** Various methods have been proposed, each with its own limitations. Some naive normal-based tests have low testing power with invalid normal distribution assumptions for RNA-seq read counts, whereas count-based methods lack a biologically meaningful interpretation and have limited capability for integration with other analysis packages for mRNA abundance. In this study, we propose an improved method, RoMA, to accurately detect differential expression and unlock the integration with upstream and downstream analyses on mRNA abundance in RNA-seq studies.

**Results:** RoMA incorporates information from both mRNA abundance and raw counts. Studies on simulated data and two real datasets showed that RoMA provides an accurate quantification of mRNA abundance and a data adjustment-tolerant DE analysis with high AUC, low FDR, and an efficient control of type I error rate. This study provides a valid strategy for mRNA abundance modeling and data analysis integration for RNA-seq studies, which will greatly facilitate the identification and interpretation of DE genes.

**Availability and implementation:** RoMA is available at https://github.com/GuoshuaiCai/RoMA.

## Background

RNA-sequencing (RNA-seq) has been widely used to quantify mRNA abundance in cells and tissues of interest. Various methods for identifying differentially expressed genes (DEG) from RNA-seq data have been proposed. One strategy is to transform read counts into continuous values representing mRNA abundance such as reads/fragments per kilobase per million mapped reads (RPKM/FPKM) (Mortazavi, et al., 2008) (Trapnell, et al., 2010), then normal-based hypothesis testing is applied to these values to identify DEG. Despite usually logarithm transformation is applied to increase the normality of the data, the assumption of normality is violated in RNA-seq data due to the inherent dependency between variance and sequencing depth, which cannot be removed by these transformations. Count-based methods have been proposed as an alternative approach to model raw read counts or counts per million (CPM) (Robinson, et al., 2010), which keep the mean-variance dependence and deal with the heteroscedasticity. Two commonly used methods, DESeq (Anders and Huber, 2010) and edgeR (Haas, et al., 2013; Robinson, et al., 2010), assume raw counts follow a negative binomial distribution and use a non-parametric local regression to capture the mean-variance dependency of the sequencing read count. Cai *et al*. modeled the proportion of read counts in two samples based on the beta binomial distribution (Cai, et al., 2012; Cai, et al., 2017). Rather than assuming a count distribution, limma-voom believed that precise capture of mean-variance dependency is most important (Law, et al., 2014). It estimated the mean-variance dependency from CPM and used it as a precision weight in the linear model to detect differentially expressed genes with compatible performance with negative binomial models (Soneson and Delorenzi, 2013).

Count-based methods have achieved high power in DE analyses and are now widely used. However, the CPM which is modeled in these methods lacks direct biological meaning and has limited ability to be integrated with many analyses on mRNA abundance. For example, ComBat models mRNA abundance by assuming bias factors affect the expression of genes in similar ways (Johnson, et al., 2007). Single sample gene set enrichment analysis (ssGSEA) works on gene rank from mRNA abundance (Barbie, et al., 2009). and MA plots and heatmaps were designed to show the relative expression abundance (Dudoit, et al., 2002). In addition, count-based DE analysis methods do not allow any transformation of counts which will break the mean-variance dependency, such as those from batch correction.

Here, we propose a new DE analysis method called <u>R</u>NA-seq analysis <u>o</u>f <u>M</u>olecular <u>A</u>bundance (RoMA), which is easy to interpret, can be integrated with analyses of mRNA abundance and provides data normalization and DE analysis. Based on limma-voom’s framework, we incorporate information from both mRNA abundance and raw counts by (1) modeling RPKM to represent the mRNA relative abundance, (2) using the mean-variance dependency from CPM as a precision weighted account of sequence depth variability and (3) moderating the t-statistic by shrinking the gene-wise variance using an empirical Bayes method (Law, et al., 2014). We developed an R toolkit (https://github.com/GuoshuaiCai/RoMA) for use by the biomedical research community.

RoMA is powerful and robust to detect DEG. RoMA tolerates data adjustment (which is inapplicable for limma-voom), such as batch correction, because of its ability to include information from both mRNA abundance and RNA-seq raw read counts. To test this merit, we simulated a scenario that RNA-seq data requires adjustment to remove the bias or effects to obtain the true biological signal. For simplicity, we simulated RNA-seq experiments with different yielding of sequencing reads due to the effects, which need to be adjusted by equaling the sequencing depth. We observed that RoMA and t-test showed the same performance on both raw data and adjustment data, whereas limma-voom was significantly affected by data adjustment although it outperforms *t*-test (Fig 1). For data without transformation, RoMA and limma-voom had similar performances on DEG detection in all simulated and real data, resulting in high AUCs, low FDRs and low type I error rates, as technically RoMA uses the same mean-variance dependency and shrinks the variance as does limma-voom. RoMA and limma-voom are significantly superior to the *t*-test in scenarios as combinations of including small/relatively large sample size and equal/unequal sequencing depth for all simulated and real datasets including MAQC datasets and TCGA breast cancer datasets.

**Figure 1.**
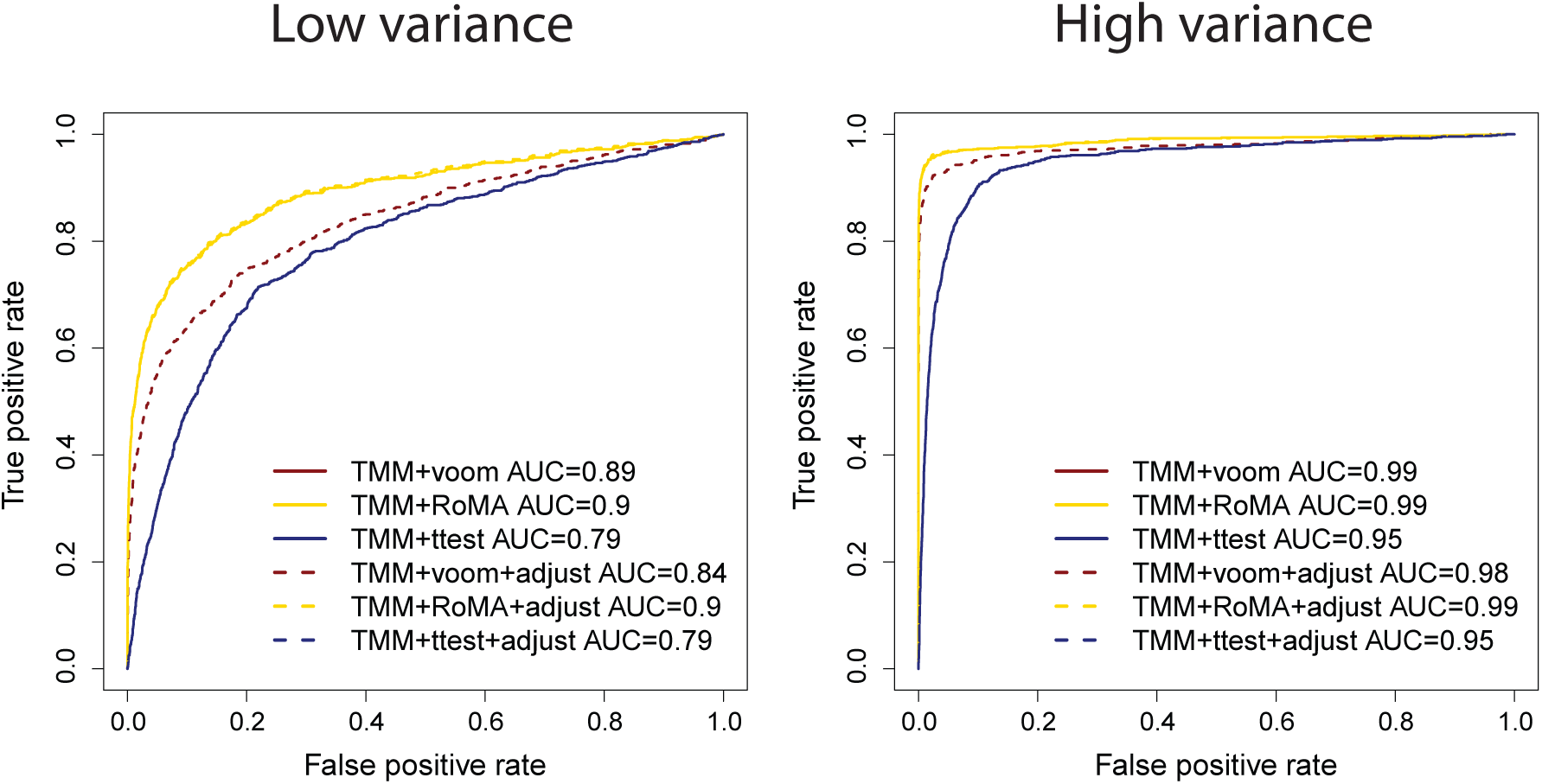
ROC curves for detecting DEG in simulated data with adjustment. TMM method was used to adjust the data to equal sequencing depths. RoMA, limma-voom and t-test were applied to perform DE analyses on both raw data and adjusted data. Red solid, yellow solid and yellow dotted lines are overlap. Blue solid and blue dotted lines are overlap. ROC curves in the left panel are shown for low variance genes with fold changes equal to 2/3 or 3/2 and those in the right panel are shown for high variance genes with fold changes equal to 1/3 or 3.

RoMA provides accurate quantification of mRNA abundance. In contrast to CPM, RoMA generates normalized RPKM representing biologically meaningful information of relative mRNA abundance, which is essential for displaying as heatmaps, MA plots, and other visualization means. RPKM/FPKM normalized data but doesn’t account for composition bias, thus we suggest using another layer of normalization to adjust it. We evaluated and benchmarked the performance of combinations of RPKM/FPKM with TMM (Robinson, et al., 2010), DESeq (Anders and Huber, 2010) and upperQuartile normalization methods (Bullard, et al., 2010) as an essential step of DEG detection. We found that TMM normalized RPKM and DESeq normalized RPKM were least biased and lead to the most precise DE detections, whereas RPKM only had the poorest performance (data not shown), which is consistent with the conclusion of the study of Dillies et al. (Dillies, et al., 2013). Therefore, we used TMM normalized RPKM as the default in RoMA.

RoMA is also effective for complicated experimental designs with multiple treatment factors and numerical covariates since it is based on linear models. By contrast, *t*-test on RPKM, which showed significantly worse performance on the control of FDR and type I error rates, should be avoided for RNA-seq DEG analyses, especially when the sample size is small.

In summary, RoMA provides robust results in terms of DEG of RNA-seq transcript abundance and a strategy for mRNA abundance modeling.

